# Basolateral amygdala to insular cortex activity makes sign-tracking behavior insensitive to outcome value

**DOI:** 10.1101/2022.02.24.481881

**Authors:** Sara E. Keefer, Daniel E. Kochli, Donna J. Calu

## Abstract

Goal-tracking rats are sensitive to Pavlovian outcome devaluation while sign-tracking rats are devaluation insensitive. During outcome devaluation, goal-tracking (GT) rats flexibly modify responding to cues based on the current value of the associated outcome. However, sign-tracking (ST) rats rigidly respond to cues regardless of the current outcome value. Our prior study demonstrated disconnection of the basolateral amygdala (BLA) and anterior insular cortex (aIC) decreased both goal- and sign-tracking behaviors. Given the role of these regions in appetitive motivation and behavioral flexibility we predicted that disrupting BLA to aIC pathway during outcome devaluation would reduce flexibility in GT rats and reduce rigid appetitive motivation in ST rats. We inhibited the BLA to aIC pathway by infusing inhibitory DREADDs (hM4Di-mcherry) or control (mCherry) virus into the BLA and implanted cannulae into the aIC to inhibit BLA terminals using intracranial injections of clozapine N-oxide (CNO). After training, we used a within subject satiety-induced outcome devaluation procedure in which we sated rats on training pellets (devalued condition) or homecage chow (valued condition). All rats received bilateral CNO infusions into the aIC prior to brief non-reinforced test sessions. Contrary to our hypothesis, BLA-IC inhibition did not interfere with devaluation sensitivity in GT rats but did make ST behaviors sensitive to devaluation. Intermediate rats showed the opposite effect, showing rigid in responding to cues with BLA-aIC pathway inactivation. Together, these results demonstrate BLA-IC projections mediate tracking-specific Pavlovian devaluation sensitivity and highlights the importance of considering individual differences in Pavlovian approach when evaluating circuitry contributions to behavioral flexibility.

## 1. Introduction

Substance Use Disorder (SUD) only affects a small portion of individuals who engage in drug use. Individuals with SUD chronically relapse into compulsive drug seeking and drug taking despite negative consequences and are less likely to change their behavior despite environmental pressures. In preclinical models, addiction vulnerability is examined with sign-tracking (ST) and goal-tracking (GT) phenotypes defined in a Pavlovian Lever Autoshaping (PLA) procedure (Flagel et al., 2009; Hearst & Jenkins, 1974). ST rats approach and vigorously engage with an insertable lever cue, a behavior that remains rigid when the associated reward is devalued. GT rats approach and engage with the food cup during the lever cue, a behavior that flexibly decreases when the associated reward is devalued (Keefer et al., 2020; Morrison et al., 2015; Nasser et al., 2015; Patitucci et al., 2016; Smedley & Smith, 2018). Here, we examine a brain pathway that is implicated in both appetitive motivation and behavioral flexibility to determine its contribution to flexibility differences in goal- and sign-tracking rats.

The basolateral amygdala (BLA) is involved in both incentive learning and motivation (Chang et al., 2012; Hatfield et al., 1996; Johnson et al., 2009; Wassum & Izquierdo, 2015), and its projections to the anterior insular cortex (aIC) are necessary for both GT and ST behaviors (Nasser et al., 2018). Contralateral disconnection of the BLA and aIC decreases goal-tracking approach and increases the latency to both goal-track and sign-track. Another study showed temporally specific involvement for the BLA and aIC during instrumental outcome devaluation, with BLA necessary for encoding the degraded outcome value and the aIC necessary for the retrieval of that outcome value at test (Parkes & Balleine, 2013). These results indicate information flow from the BLA to aIC is necessary in behavioral flexibility. Similarly, communication between the BLA and the orbitofrontal cortex (OFC), which borders the aIC, is critical for behavioral flexibility across species (Fiuzat et al., 2017; Baxter et al., 2000) and direct BLA to OFC projections are necessary for Pavlovian, but not instrumental, outcome devaluation (Lichtenberg et al., 2017). These findings indicate communication from the BLA to aIC is necessary for GT behaviors and for behavioral flexibility in instrumental outcome devaluation (for review, see Keefer et al., 2021).

The current study first examines if communication from the BLA to aIC is necessary for Pavlovian outcome-specific satiety devaluation. Then, we wanted to determine the extent to which tracking-specific pathway utilization mediates goal-and sign-tracking differences in devaluation sensitivity. We hypothesized that intact GT rats would be devaluation sensitive and that chemogenetic inhibition of the BLA to aIC pathway would make GT rats devaluation insensitive (Keefer et al., 2020; Kochli et al., 2020; Nasser et al., 2015). Furthermore, we hypothesized that intact ST rats would be devaluation insensitive to and that chemogenetic inhibition of the BLA to aIC pathway would generally reduce sign-tracking (Nasser et al., 2018) or potentially make them devaluation sensitive (Keefer et al., 2020; Kochli et al., 2020; Nasser et al., 2015). To inactivate the direct pathway from BLA to aIC, we expressed inhibitory chemogenetic constructs into bilateral BLA and implanted bilateral guide cannulae into the aIC to directly inhibit BLA terminals in aIC during outcome-specific satiety devaluation.

## 2. Materials and Methods

### 2.1 Subjects

Male and female Long-Evans rats (Charles River Laboratories, Wilmington, MA, USA; approximately 8 weeks of age upon arrival; N = 160 run as 5 cohorts) were maintained on a 12 hr light/dark cycle with lights off at 0900. Rats were doubled-housed upon arrival with *ad libitum* access to standard laboratory chow and water, and single-housed housed after acclimation and prior to surgery or behavioral procedures. We surgerized two cohorts of rats prior to all behavioral training and testing and surgerized three cohorts after determining tracking phenotype but before devaluation testing. (Results were similar regardless of surgery and behavioral timeline). We performed all behavioral procedures during the dark phase on the cycle. During all behavioral training and testing, we food-restricted rats to ∼90% of their maximum achieved body weight. We conducted all experiments in accordance to the “Guide for the Care and Use of Laboratory Animals” (8^th^ edition, 2011, US National Research Council) and were approved by University of Maryland, School of Medicine Institutional Animal Care and Use Committee (IACUC).

## 2.2 Surgical Procedures

We anesthetized rats with isoflurane (Vetone, Boise, ID, USA; 5% induction, 1-3% maintenance throughout surgery). We placed rats in a stereotaxic apparatus (model 900, David Kopf Instruments, Tujunga, CA, USA) and maintained rats body temperature with a heating pad throughout surgery. We administered subcutaneous injection of carprofen analgesic (5 mg/kg) and a subdermal injection of the local anesthetic lidocaine (10 mg/ml) at the incision site prior to first incision. We leveled the skull based on Bregma and lambda dorsal-ventral plane and performed craniotomies above each injection site with a drill. We used a 10 μl Hamilton syringe (Hamilton, Reno, NV, USA) to deliver the virus into bilateral BLA using the following coordinates: AP -2.8 mm, ML ± 5.0 mm, DV -8.5 mm 0° from midline relative to Bregma surface. We infused 600 nl of AAV8-hSyn-hM4Di-mCherry (hM4Di) or AAV8-hSyn-mCherry (mCherry; Addgene, Watertown, MA, USA) into each BLA via a micropump (UltraMicroPump III, World Precision Instruments, Sarasota, FL, USA) at a rate of 250 ml/min and the syringe was left in place for 10 min post infusion to allow for viral diffusion. After closing BLA craniotomies with bone wax, we implanted 23-gauge guide cannulae bilaterally 1 mm above our target region in the aIC using the following coordinates: AP +2.8 mm, ML ± 4.0 mm, DV -4.8 mm 0° from midline relative to Bregma surface. We anchored cannulae with jeweler’s screws and dental cement and inserted obturators into the guide cannula, which were removed periodically throughout recovery and training to ensure patency. We moved rats to a recovery cage on a heating pad, administered carprofen analgesic (5 mg/kg s.c.) at 24 and 48 hr post-surgery, and monitored their health until behavioral procedures.

### 2.3 Apparatus

We conducted behavioral experiments in identical behavioral chambers (25 × 27 × 30 cm; Med Associates) located in a different room than the colony room. Each chamber was contained in individual sound-attenuating cubicles with a ventilation fan. On one wall, a red house light (6 W) was illuminated during PLA sessions and devaluation tests. The opposite wall had a receded food cup with photobeam detectors, and the cup was 2 cm above the grid flood. A programmed pellet dispenser was attached to the food cup and delivered 45 mg food pellets (catalog # 1811155; Test Diet Purified Rodent Tablet [5TUL]; protein 20.6%, fat 12.7%, carbohydrate 66.7%). One retractable lever was located 6 cm above the grid floor on either side of the food cup, and lever side was counterbalanced between subjects.

### 2.4 Pavlovian Lever Autoshaping

Prior to training, we habituated rats to the food pellets in their home cage to reduce novelty to the food. Next, we trained rats on daily Pavlovian lever autoshaping (PLA) sessions that lasted ∼26 min and included 25 presentations of non-contingent lever presentations (conditioned stimulus; CS) and occurred on a VI 60 s schedule (50-70 s). After the 10 s lever presentation, the lever retracted and two 45 mg food pellets were delivered via food cup. After 25 trials, the red house light turned off, and we returned rats to their cage and colony room.

### 2.5 Satiety-Induced Outcome Devaluation Testing

After the 5^th^ PLA session, we gave rats two sessions of satiety-induced outcome devaluation testing. Rats had one hour of access to 30 g of either their homecage chow (valued condition) or food pellets used during PLA training (devalued condition) in a pre-habituated ceramic ramekin. Within 15 min of the end of satiation hour, we infused 0.25 μl of 1 mM clozapine N-oxide (CNO; Tocris or Abcam) dissolved in aCSF into bilateral aIC over 1 min and left injectors in place for an additional minute to allow to diffusion of solution. We waited 10-15 min after infusion to allow binding of the CNO to the DREADD receptors, and then placed rats into the behavioral chambers for devaluation probe test. Tests consisted of 10 non-rewarded lever presentations on VI 60s schedule (50-70 s). After test, we gave rats a 30 min food choice test in their homecage, which included 10 g of chow and 10 g of food pellets in separate ramekins to confirm satiety was specific to the outcome they were pre-fed.

### 2.6 Behavioral Measurements

For PLA training sessions and devaluation probe tests, we recorded number and duration of contacts, latency to contact, and probability of contact for each behavior to the food cup and the lever during the 10 s CS (lever) period. On trials with no contact, a latency of 10s was recorded. Probability of contact was calculated by determining the number of trials that the lever or food cup contact was made, divided by total number of trials in that session.

To determine tracking phenotype, we used a Pavlovian Conditioned Approach (PCA) analysis (Meyer et al., 2012) which quantifies the continuum of lever-directed (sign-tracking) and food cup-directed (goal-tracking) behaviors. PCA scores are the average of three separate score measures: (1) preference score, (2) latency score, and (3) probability score. Preference score is number of lever contacts minus number of food cup contacts during the CS divided by the sum of these two measures. Latency score is the average time to make a food cup contact minus the time to make a lever contact during the CS divided by 10 s (the duration of the CS). Probability score is the probability to make a lever contact minus probability to make a food cup contact across trials in a session. PCA score for each rat was determined by averaging the PCA scores for PLA sessions 4 and 5. Sign-tracking PCA range from +0.25 to +1.0, goal-tracking PCA range from -0.25 to -1.0, and intermediate PCA range from -0.24 to +0.24.

For devaluation probe tests, we examined total behavior (sum of food cup and lever contacts during the 10 s CS period) and responding of each behavior separately. We also examined preferred responding – contacts to food cup for goal-trackers and contacts to lever for sign-trackers. For consumption data on test days, we recorded the amount of pellets and chow in grams during satiety hour and during the 30 min choice test.

### 2.7 Histology

After all behavioral training and testing finished, we anesthetized rats with isoflurane and transcardially perfused with 100 ml of 0.1 M PBS then 400 ml of 4% paraformaldehyde in 0.1 M sodium phosphate, pH 7.4. We extracted brains and post-fixed them in the 4% paraformaldehyde solution for at least 2 hr prior to incubation in a 30% sucrose in 0.1 M sodium phosphate for at least 24 hr at 4°C. We rapidly froze brains in dry ice and stored them in -20°C until slicing. Using a cryostat (Leica Microsystems), we collected 30 μm sections into 4 series through the cannulae placement in aIC and through the virus infusion sections in the BLA. Sliced tissue was stored in cryopreservant in -20°C until mounting or immunohistochemistry. We mounted cannulated aIC sections onto gelatin-coated slides, and after drying, we stained with cresyl violet, coverslipped with Permount, and examined under a light microscope for confirmation of cannulae placement into the aIC. We mounted BLA sections onto SuperFrost slides, and after drying, we coverslipped with Vectashield mounting medium with DAPI. We used immunohistochemistry to amplify hM4Di-mCherry expression on the terminals in the aIC for confirmation of terminal expression. Floating aIC sections were rinsed in 0.1 M PBS 2 times for 10 min and blocked for 1 hr with 2% normal goat serum and 0.3% Triton X-100 in PBS. Sections were rinsed in PBS twice for 10 min and incubated in blocking solution with anti-dsRed primary antibody raised in rabbit (1:500; Clontech 632496) overnight in 4°C with gentle agitation. Sections were rinsed 2 times for 10 min each in the blocking solution, then incubated in blocking solution containing AlexaFluor-594 goat anti-rabbit (1:500; Invitrogen). After three 10 min rinses in PBS, we mounted sections onto SuperFrost slides and coverslipped with Vectashield mounting medium with DAPI. We confirmed viral expression in the BLA and in terminals within the aIC under 5x or 10x using a Confocal SP8 (Leica Microsystems) and used anatomical boundaries defined by (Paxinos & Watson, 2007; Swanson, 2004). We excluded rats if cannulae or viral placements were outside the region of interest (Fig. 4), or if terminal expression could not be confirmed, which resulted in N=75 (37 females, 38 males): GT: 13 mCherry, 12 hM4Di; ST: 13 mCherry, 13 hM4Di; and INT: 13 mCherry, 11 hM4Di.

### 2.8 Statistical Analysis

We analyzed data using SPSS statistical software (IBM v.25). We used mixed-design repeated measures ANOVAs. When applicable, the within-subject factors were Response (food cup, lever) and Outcome Value (valued, devalued), and the between-subject factors were Virus (hM4Di, mCherry), Tracking Group (ST, INT, GT), and Sex (female, male). Significant main effects and interactions were followed by post-hoc paired samples or independent t-tests.

## 3. Results

### 3.1 Limited Pavlovian Lever Autoshaping

Prior to devaluation testing, we trained rats on five sessions of Pavlovian Lever Autoshaping (PLA) to determine tracking phenotype by examining lever- and food-cup directed behaviors. Tracking phenotype is determined by a rat’s Pavlovian Conditioned Approach Index (Fig. 1A; see methods for calculation) on the last two sessions of PLA prior to devaluation testing and is based on the difference between the number of lever presses (Fig. 1B) and food cup pokes (Fig. 1C) as well as the difference score for latency and probability to engage with the lever and food cup. In Table 1, we report the main effects and interactions for the autoshaping data using six separate mixed-design repeated measures ANOVAs. Tracking group (ST, INT, GT) was the between-subjects factor, and Session (1-5) was the within-subjects factor. To confirm there were no differences between viral groups (mCherry, hM4Di) within each tracking group (ST, INT, GT) prior to devaluation testing, we analyzed number of lever and food cup contacts during the 5^th^ session of PLA. We found no main effects of Virus nor Virus x Tracking Group interactions (*F*s < 2.89, *p*s > 0.05; Fig. 1D).

**Table 1.** Repeated measures analysis of variance (ANOVA) for Pavlovian lever autoshaping across all tracking groups during limited training (sessions 1-5).

| Effect | Degrees of Freedom | Contact |  | Latency |  | Probability |  |
| --- | --- | --- | --- | --- | --- | --- | --- |
|  |  | *F* | *p* | *F* | *p* | *F* | *p* |
| Lever Presses |  |  |  |  |  |  |  |
| Session | (4,288) | 83.92 | <0.001 | 144.89 | <0.001 | 122.18 | <0.001 |
| Tracking Group | (2,72) | 73.03 | <0.001 | 84.35 | <0.001 | 88.50 | <0.001 |
| Session x Tracking group | (8,288) | 21.36 | <0.001 | 26.93 | <0.001 | 19.00 | <0.001 |
| Food cup pokes |  |  |  |  |  |  |  |
| Session | (4,288) | 24.14 | <0.001 | 37.98 | <0.001 | 25.01 | <0.001 |
| Tracking Group | (2,72) | 35.26 | <0.001 | 51.09 | <0.001 | 39.81 | <0.001 |
| Session x Tracking group | (8,288) | 19.35 | <0.001 | 18.34 | <0.001 | 12.48 | <0.001 |

**Figure 1.**
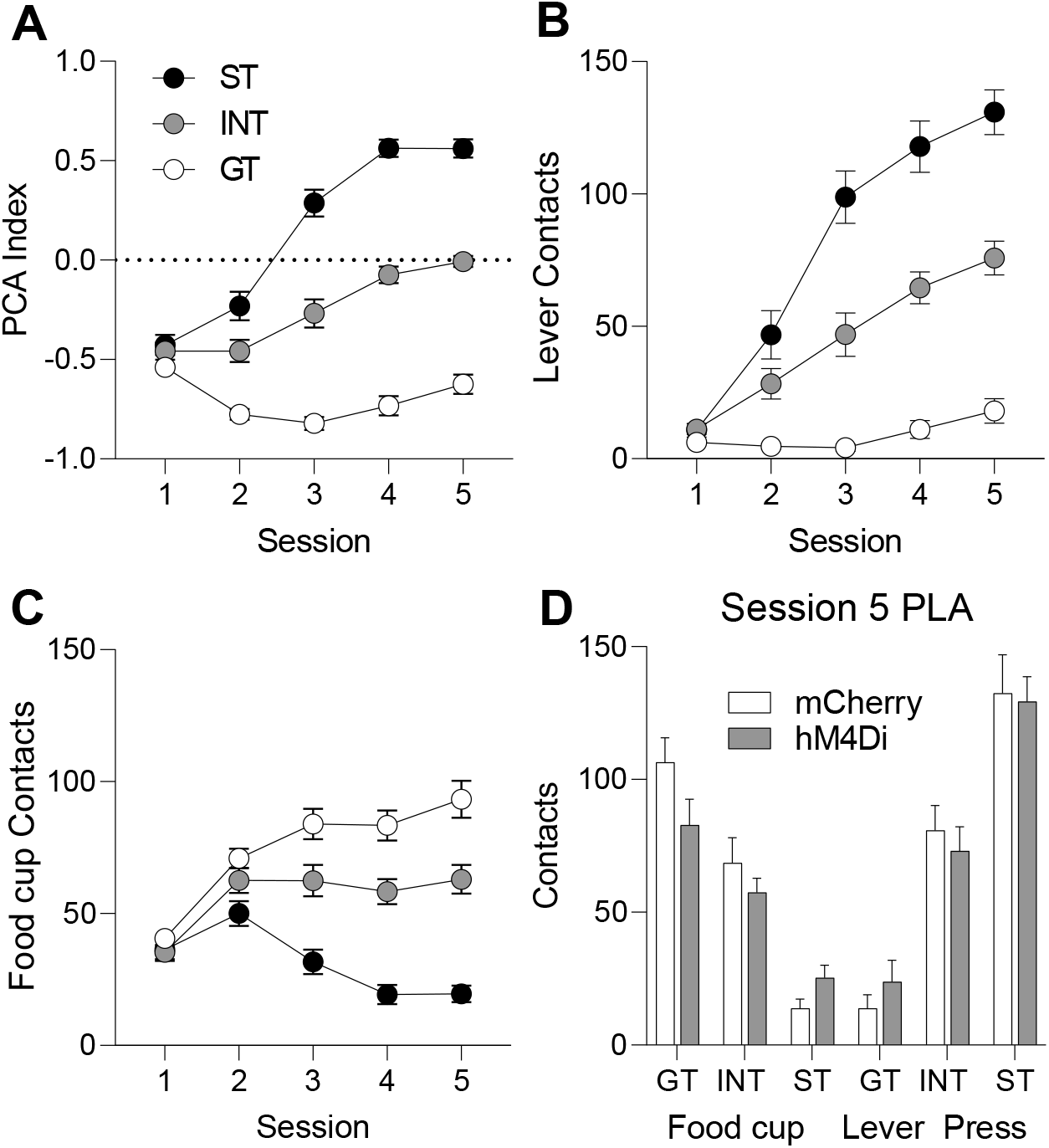
Pavlovian Lever Autoshaping (PLA) data. **A)** Pavlovian Conditioned Approach (PCA) index, **B)** lever contacts, and **C)** food cup contacts across five days of training. **D)** Lever and food cup contacts on the fifth day of PLA across tracking groups and between viral conditions. There were no differences between viral conditions within each tracking group (*p*s > 0.05). Data are mean ± SEM.

### 3.2 Satiety-Induced Outcome Devaluation After Limited Training

We examined if inactivation of BLA-aIC altered Pavlovian outcome devaluation independent of tracking group (Fig. 2A). We first analyzed total behavior (sum of lever and food cup contacts) using the between subjects factor of Virus (mCherry, hM4Di) and within subjects factor of Devaluation (Valued, Devalued). We observe a main effect of Devaluation (*F*(1,73) = 22.67, *p* < 0.001) but no other main effects or interactions (*F*s < 0.2, *p*s > 0.5). Next, we repeated the same analysis separately on lever (Fig. 2B) and food cup (Fig. 2C) behaviors, and found a similar result: main effect of Devaluation (lever contacts: *F*(1,73) = 10.34, *p* < 0.01; food cup contacts: *F*(1,73) = 24.94, *p*<0.001) but no other main effects or interactions (*F*s < 0.6, *p*s > 0.4). Without considering tracking differences in Pavlovian approach, it appears as though BLA-IC pathway inhibition has no effect on responding to cues when outcome value changes.

**Figure 2.**
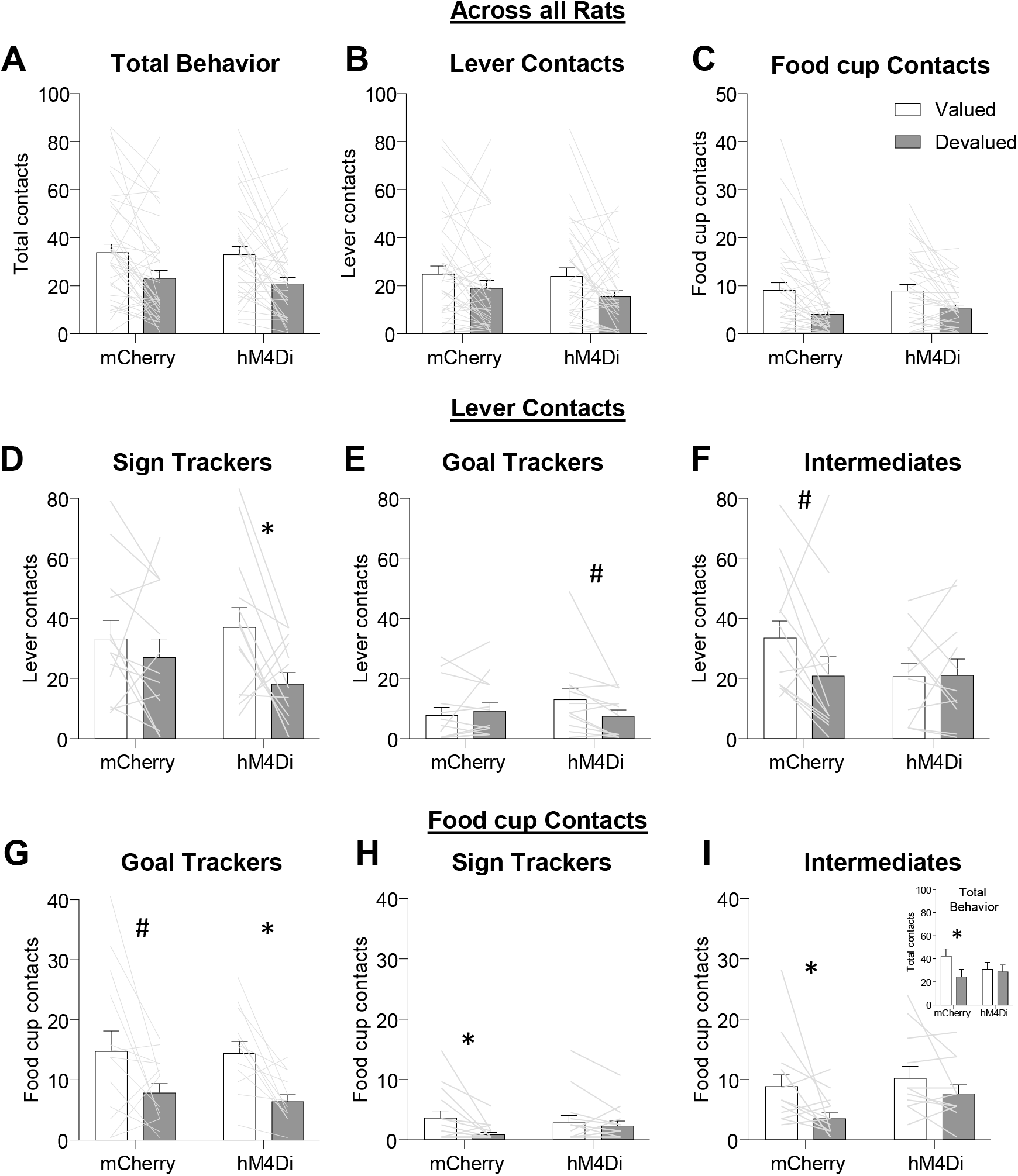
Specific satiety-induced outcome devaluation in sign-tracking, goal-tracking, and intermediate rats. Data are represented as within-subject individual data (lines) and group averages (bars; mean + SEM). Overall main effects of Outcome Value on **A)** total behavior (lever + food cup contacts) and separately on lever contacts **B)** and food cup contacts **C)**, with no main effects of Virus nor interaction (*p*s > 0.05). **D-F)** For lever contacts, we observe a Virus x Tracking x Outcome Value interaction (see Results), and *post hoc* analysis indicate intact (mCherry) ST and GT show devaluation insensitivity, while intact (mCherry) INT are marginally devaluation sensitive (*t*(12) = 2.07, *p* = 0.061). While ST and GT rats with BLA-aIC inhibition were devaluation sensitive (ST hM4Di: *t*(12) = 2.63, *p* = 0.022; GT hM4Di: *t*(11) = 2.12, *p* = 0.057), INT rats with BLA-Aic inhibition were devaluation insensitive (INT hM4Di: *t*(10) = -0.10, *p* = 0.926). **G-I)** For food cup contacts we observe a main effect of Outcome Value, and our *a priori* planned comparisons confirm devaluation sensitivity in both GT viral groups (GT mCherry marginal: *t*(12) = 2.08, *p* = 0.059; GT hM4Di *t*(12) = 3.94, *p* = 0.002). See Results for ST and INT food cup data reporting. **I, inset**) Total behavior (sum lever and food cup contacts for intermediate rats). Planned comparisons show intact INT are devaluation sensitive (INT mCherry: *t*(12) = 2.59, *p* = 0.024) but INT rats with BLA-aIC inhibition were devaluation insensitive (INT hM4Di *t*(10) = 0.359, *p* = 0.727). * *p* < 0.05, # *p* = 0.06.

However, we designed the study to determine if tracking groups uniquely utilized the BLA-aIC pathway to drive their differential devaluation sensitivities. Thus, we included tracking phenotype as a between subjects factor and separately analyzed lever and food cup contacts, the dominant behaviors of sign- and goal-trackers, respectively. For lever contacts (Fig. 2D-F), we observe a Virus x Tracking Group x Outcome Value interaction (*F*(2,69) = 3.28, *p* = 0.044) and main effects of Tracking Group (*F*(2,69) = 12,12, *p* < 0.001) and Outcome Value (*F*(1,69) = 10.45, *p* = 0.002), but no main effect of Virus or any other interaction (*F*s < 2.2, *p*s > 0.1). *Post hoc* analyses indicate that intact ST mCherry rats show no difference in lever contacts to valued and devalued conditions (Fig. 2D; *t*(12) = 1.05, *p* = 0.316), however, ST hM4Di expressing rats show greater lever approach to the valued compared to devalued condition (*t*(12) = 2.63, *p* = 0.022). While goal-tracking rats show very low levels of lever approach (i.e. sign-tracking behavior), they showed a similar pattern of results with BLA-aIC inactivation (Fig. 2E; GT mCherry: *t*(12) = -0.75, *p* = 0.466; GT hM4Di: *t*(11) = 2.12, *p* = 0.057). To our surprise, intact INT mCherry rats showed marginally greater lever contacts to valued compared to devalued conditions (Fig. 2F; *t*(12) = 2.07, *p* = 0.061) and pathway inactivation in INT hM4Di rats made lever contacts devaluation insensitive (*t*(10) = -0.10, *p* = 0.926). These data suggest that for the extreme ends of the tracking continuum (ST and GT rats), disrupting communication between BLA and aIC makes lever approach (i.e. sign-tracking behavior) more sensitive to current outcome value. In contrast, rats displaying a mix of lever and food cup approach (INT rats), BLA-aIC inactivation makes lever approach less sensitive to current outcome value. Indeed, an analysis that combines ST and GT lever contact data (i.e. the PCA continuum extremes, STGT group) and compares it to lever data from intermediates supports these conclusions, with a Tracking (STGT, INT) x Virus x Outcome Value interaction (*F*(1,71) = 6.187, *p* = 0.015).

For food cup contacts (Fig. 2G-I), we observe a main effect of Outcome Value (*F*(1,69) = 27.03, *p* < 0.001) and Tracking Group (*F*(2,69) = 21.53, *p* < 0.001) and Outcome Value x Tracking Group interaction (*F*(2,69) = 4.20, *p* = 0.019). While we did not observe a three-way interaction for food cup contacts, our *a priori* hypothesis was that the preferred response (i.e. food cup contact) in goal-tracking rats would become devaluation insensitive with BLA-aIC pathway inhibition. Contrary to our predictions, both GT mCherry and hM4Di rats showed more food cup contacts to valued compared to devalued conditions (Fig. 2G; main effect of Outcome Value (*F*(1,23) = 14.16, *p* < 0.01). While the remaining food cup approach data should be interpreted with caution due to low levels or responding and/or a lack of three-way interaction, we also observe a main effect of Outcome Value in both ST (Fig. 2H; *F*(1,24) = 6.19, *p* = 0.02) and INT groups (Fig. 2I; *F*(1,22) = 7.56, *p* = 0.012).

Altogether, the data across tracking groups suggests BLA-aIC inhibition does not affect the devaluation sensitivity of food cup approach. Because INT rats display similar levels of food cup and lever approach, Fig. 2I inset shows total approach (lever + food cup) for INT rats.

Because we used both males and females in this study we also analyzed the data using Sex instead of Tracking as a factor. The ANOVA including between subject factors of Sex and Virus and within subject factor of Response and Outcome Value yielded main effect of Sex (*F*(1,71) = 11.93, *p* < 0.001), and a Response x Sex interaction (*F*(1,71) = 5.95, *p* = 0.017), but no other main effects or interactions. Consistent with prior studies, we observe greater levels of lever approach in females compared to males (Fig. 3A; *t*(73) = 3.077, *p* = 0.003).

**Figure 3.**
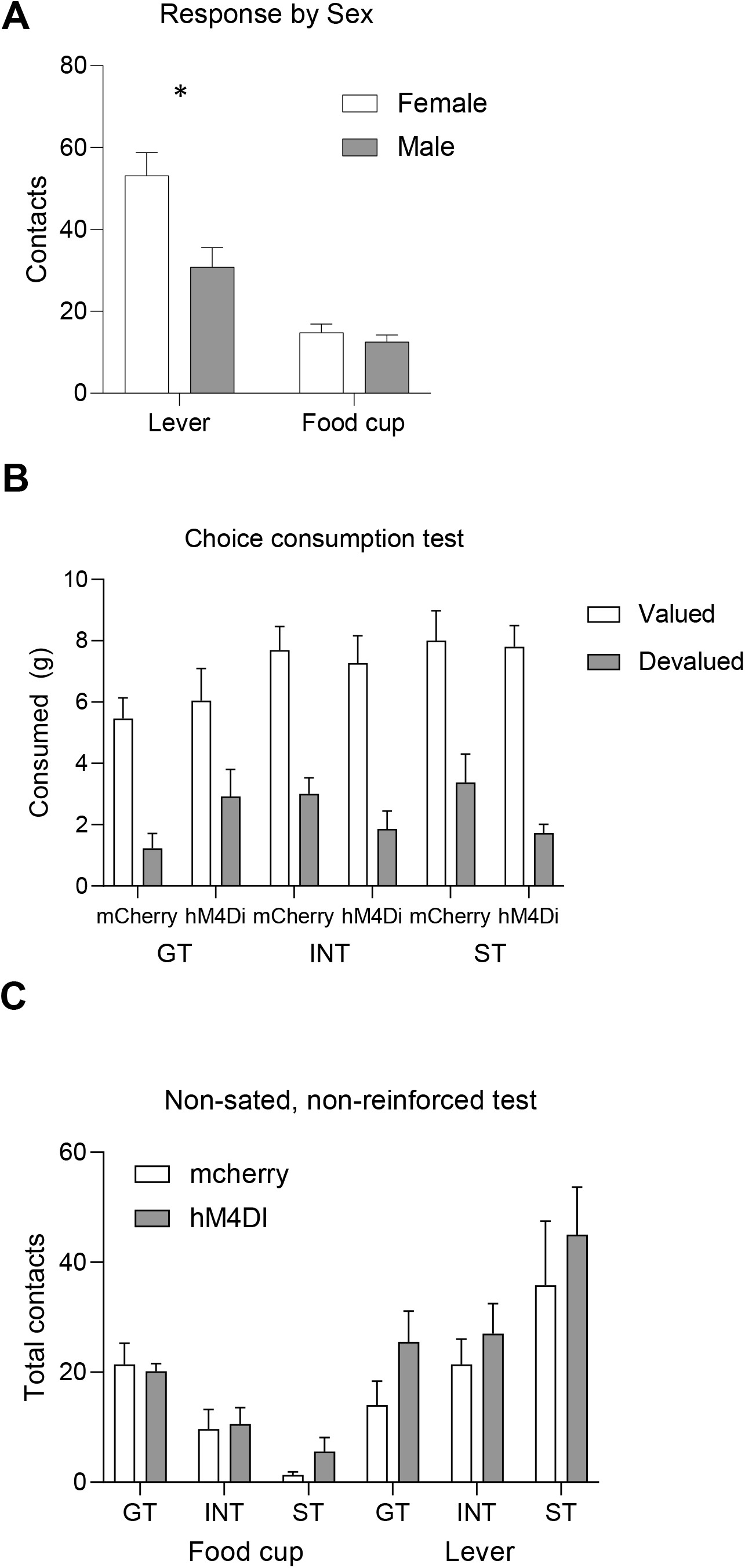
**A)** During outcome devaluation, we observed a Response x Sex interaction, and *post hoc* analysis show females perform more lever contacts (ie. sign-tracking responses) than males during outcome devaluation (t(73) = 3.077, *p* = 0.003), independent of Outcome Value or Viral condition. There are no differences in food cup contacts. **B)** We found no differences between tracking or viral groups during the post-outcome devaluation consumption choice test, indicating neither tracking nor BLA-aIC inhibition affected choice to consume the valued over the devalued outcome. **C)** We found no effects of BLA-aIC inactivation on food cup or lever contacts during a non-sated, non-reinforced test.

**Figure 4.**
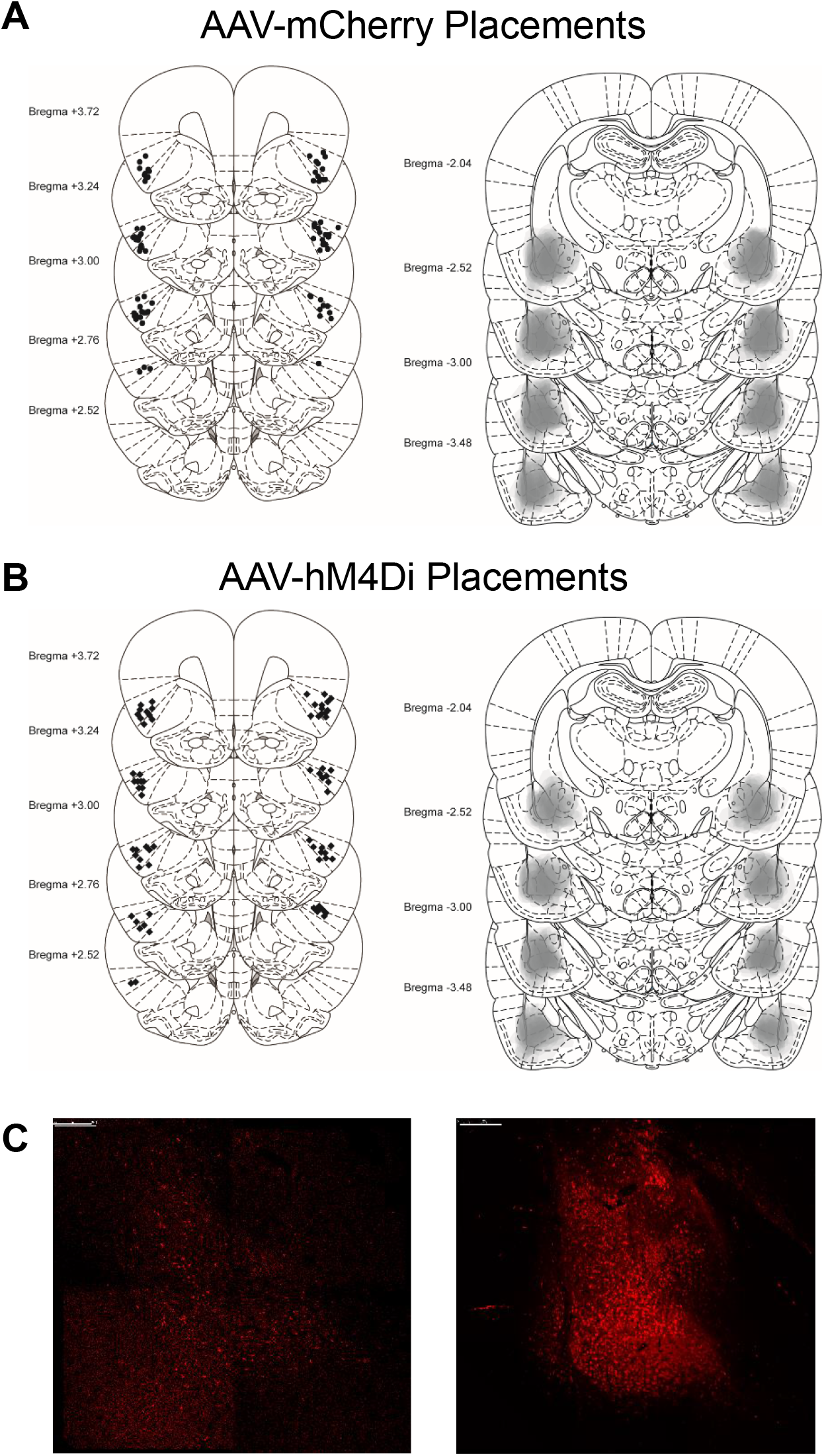
Histological verification of cannulae in the aIC and viral expression in the BLA. We implanted bilateral aIC cannulae (**A**, left; **B**, left) and infused viral constructs into the BLA (**A** right, **B** right; Paxinos and Watson, 2007). Representative images of hM4Di (**C**, left), and mCherry (**C**, right). Scale bar = 250 μm.

### 3.3 Consumption, Choice Test, and Non-sated probe test

We sated rats on either chow (Valued) or pellets (Devalued) prior to devaluation probe test. We found no differences in the amount of food consumed between Tracking Groups or Virus conditions (*F*s < 2.9, *p*s > 0.09). To confirm devaluation of the sated food, we gave rats a choice test between the food they were sated on and the other food. Rats consumed less of the food they were sated on and more of the alternative food (Fig. 3B), verified by a main effect of Choice (*F*(1,69) = 169.38, *p* < 0.001), with no main effects of Tracking or Virus (*F*s < 2.6, *p*s > 0.07) and no interactions (*F*s < 2.14, *p*s > 0.1).

To examine if inactivation of BLA-aIC altered lever or food cup approach independent of specific satiety, we conducted a non-sated, non-reinforced test. In a mixed ANOVA with between subject factors of Virus (mCherry, hM4Di) and Tracking (ST, INT, GT) and within subjects factor of Response (lever, food cup), there was no main effect of Virus nor interactions with Virus (*F*s < 2.5, *p*s > 0.1; Fig. 3C), suggesting that BLA-aIC inactivation did not affect lever or food cup behaviors when rats were not sated.

## 4. Discussion

In the current study, we examined if communication from the BLA to the aIC is necessary for Pavlovian specific satiety-induced outcome devaluation. While we did not observe overall effect of the manipulation on outcome devaluation, when we include tracking phenotype in the analysis, we observe tracking specific effects of BLA-aIC pathway inhibition on devaluation sensitivity. Consistent with previous findings, we find that food cup behavior of intact GT rats is devaluation sensitive while lever-directed behavior of intact ST rats is devaluation insensitive (Keefer et al., 2020; Kochli et al., 2020; Nasser et al., 2015; Patitucci et al., 2016; Smedley & Smith, 2018). For the sign-tracking response (i.e. lever-directed behaviors), BLA-aIC inhibition promoted devaluation sensitivity in ST (and to small extent in GT) rats, but devaluation insensitivity in INT rats. BLA-aIC inhibition had minimal effect on devaluation sensitivity of the goal-tracking response (i.e. food cup directed behaviors). Yet the qualitatively consistent effects of BLA-aIC inhibition on both lever and food cup behavior of INT rats suggests the BLA-aIC pathway may be promoting behavioral flexibility in these rats, while the same pathway supports rigid sign-tracking behaviors for rats on extreme ends of the tracking continuum.

Previously, we observed that BLA-aIC communication is necessary for full expression of sign- and goal-tracking behaviors (Nasser et al., 2018). Contralateral disconnection of the BLA and aIC with baclofen/muscimol decreased food cup approach (in GT rats) and increased the latency to contact both the food cup (in GT rats) and lever (in ST rats), seemingly disrupting both goal- and sign-tracking behaviors. Based on our findings that goal-tracking rats are devaluation sensitive but sign-tracking rats are not, we hypothesized that the content of the associative information encoded in BLA-aIC pathway may differ between sign- and goal-tracking rats. We predicted that goal-tracking rats use BLA-aIC to encode flexible stimulus-outcome associations that are necessary for outcome devaluation sensitivity (Holland, 1998). We predicted that sign-tracking rats use BLA-aIC to encode rigid stimulus-response associations that are insensitive to outcome devaluation (Holland & Rescorla, 1975). Qualitatively, the effect of BLA-aIC inactivation on Pavlovian approach that we hypothesized for GT rats, we observed for INT rats. While intermediate rats display both lever and food cup directed behaviors, intermediate to that of sign- and goal-tracking rats, their behavior with BLA-aIC pathway inactivation is far from intermediate to the effects of this pathway manipulation on sign- and goal-tracking behaviors. We interpret the behavior of intermediates in the context of other studies below.

Consistent with our predictions for ST rats, we do find evidence for rigid encoding of stimulus-response associations in the BLA-aIC pathway, as inhibiting this pathway makes ST rats sensitive to outcome devaluation. Both the BLA and aIC are heavily implicated in appetitive motivational processes (see Centanni et al., 2021; Izquierdo, 2017; Parkes et al., 2018). The BLA is necessary for incentive processes, particularly for the expression of sign-tracking (Chang et al., 2012), and other incentive learning processes, such as second-order conditioning (Hatfield et al., 1996; P. C. Holland, 2016; Setlow et al., 2002) and conditioned reinforcement (Burke et al., 2007; Parkinson et al., 2001), for reviews, see Keefer et al., 2021; Wassum & Izquierdo, 2015). This finding is consistent with our prior study reporting that inhibition of BLA-nucleus accumbens core communication also makes sign-tracking rats sensitive to outcome devaluation (Kochli et al., 2020). Together our studies suggest an overabundance of rigid appetitive encoding in BLA projections makes sign-tracking rats insensitive to outcome devaluation (Kochli et al., 2020; Nasser et al., 2015).

Perhaps most surprising is our failure to observe effects of BLA-aIC pathway inactivation in GT rats; if anything, GT rats expressing the inhibitory DREADD construct showed qualitatively stronger devaluation than intact rats, perhaps suggesting that encoding in BLA-aIC may also support rigid stimulus-response associations in GT rats, similar to what we observe in ST rats. We consider these findings in the context of a previous study that showed temporally specific engagement of the BLA and IC during specific satiety outcome devaluation. In a series of experiments, Parkes and Balleine (2013) demonstrated the BLA is necessary for updating outcome value during satiation, but not necessary for the retrieval of this new value during test, findings consistent with BLA’s role in Pavlovian devaluation (Hatfield et al., 1996; Johnson et al., 2009; Setlow et al., 2002; West et al., 2012). Then, they demonstrated the IC is necessary for the retrieval of the new outcome value at test, but not for the initial encoding of the outcome value during satiety (Parkes et al., 2018; Parkes & Balleine, 2013). Since we sated rats prior to BLA-aIC inhibition, one possible explanation for lack of effects in GT rats is that the BLA may have already updated aIC on the new outcome value so that it was successfully retrieved at test to support devaluation sensitivity (see Piette et al., 2012). Inhibition of the BLA-aIC pathway prior to satiation may result in outcome devaluation insensitivity in otherwise devaluation sensitive behaviors, an avenue for future research. Interpreting the improvement in flexibility in ST rats with BLA-aIC inactivation within this framework suggests the rigid stimulus-response association persist beyond the encoding stage (satiety) and is dominant at time of retrieval. The opposite appears to be the case in INT rats, in which BLA-aIC communication of updated outcome value (ie. flexible stimulus-outcome association) is necessary for devaluation sensitivity at time of memory retrieval.

Unlike our prior study, disrupting communication between BLA and aIC did not disrupt sign- and goal-tracking. Our prior BLA-aIC inactivation study (Nasser et al., 2018) disrupted bidirectional communication between the BLA and aIC during a reinforced lever autoshaping test. In contrast, the current test was under extinction conditions and inhibited direct communication from BLA to aIC, while leaving communication from the aIC to BLA intact. The reinforced versus extinction conditions may account for the difference. However, it may be the case that our prior effects of contralateral inactivation on goal-tracking could be due to aIC to BLA communication. In support of this hypothesis, several studies have probed the necessity of the neighboring orbitofrontal cortex (OFC), which also lesioned the aIC. These studies concluded the OFC/aIC is necessary to retrieve and express the value of the outcome and associated cues during periods of behavioral flexibility (e.g. Gallagher et al., 1999; Ostlund & Balleine, 2007; Pickens et al., 2003, 2005). Additional studies indicate that communication between the BLA and OFC are necessary for outcome devaluation (Baxter et al., 2000; Fiuzat et al., 2017), and direct projections from the BLA to OFC and from the OFC to BLA are critical for Pavlovian, but not instrumental, outcome devaluation (Lichtenberg et al., 2017; Malvaez et al., 2019). Similarly, communication between the BLA and OFC is necessary for other behavioral flexibility paradigms, such as over-expectation (Lucantonio et al., 2015), outcome-specific Pavlovian-to-Instrumental transfer (Lichtenberg et al., 2017; Sias et al., 2021), and risky decision making (Zeeb & Winstanley, 2013; see Keefer et al., 2021 for review).

Altogether, we conclude by suggesting the utility of the Pavlovian Lever Autoshaping procedure for identifying individual differences that elucidate unique pathway contributions to behavioral flexibility.

## Acknowledgements

NIDA R01DA043533 to DJC, NIDA F32DA053772-01 to SEK, McKnight Memory and Cognitive Disorders Award to DJC (McKnight Foundation), Brain and Behavior Research Foundation NARSAD Young Investigator Grant # 24950 to DJC. Daniel E. Kochli’s current affiliation is Washington College, Psychology Department, 300 Washington Avenue, Chestertown, MD 21620, USA.

